# Controlling morpho-electrophysiological variability of neurons with detailed biophysical models

**DOI:** 10.1101/2023.04.06.535923

**Authors:** Alexis Arnaudon, Maria Reva, Mickael Zbili, Henry Markram, Werner Van Geit, Lida Kanari

**Affiliations:** Blue Brain Project, École polytechnique fédérale de Lausanne (EPFL), Campus Biotech, Geneva, Switzerland

## Abstract

Variability is a universal feature among biological units such as neuronal cells as they enable a robust encoding of a high volume of information in neuronal circuits and prevent hyper synchronizations such as epileptic seizures. While most computational studies on electrophysiological variability in neuronal circuits were done with simplified neuron models, we instead focus on the variability of detailed biophysical models of neurons. With measures of experimental variability, we leverage a Markov chain Monte Carlo method to generate populations of electrical models able to reproduce the variability from sets of experimental recordings. By matching input resistances of soma and axon initial segments with the one of dendrites, we produce a compatible set of morphologies and electrical models that faithfully represent a given morpho-electrical type. We demonstrate our approach on layer 5 pyramidal cells with continuous adapting firing type and show that morphological variability is insufficient to reproduce electrical variability. Overall, this approach provides a strong statistical basis to create detailed models of neurons with controlled variability.

## Introduction

Neurons in the brain are highly heterogeneous, both in terms of morphologies and electrical phenotypes. Attempts have been made to classify them into morphological, electrical, or combined morpho-electrical types [Gupta et al., 2000, pet, 2008, Gouwens et al.,2019], however even within a single cell type, cells are highly variable. For example, morphologies grow in the available space following the local signalling processes, hence their shapes and sizes are location-specific and unique [Galloni et al., 2020] and have an impact on electrical properties [Eyal et al., 2014]. Morphological variability has been studied in the context of robust circuit structural connectivity for example in [Hill et al., 2012, Ramaswamy et al., 2012, Croxson et al., 2018]. Also, the electrophysiological features of cells within a firing type are often highly variable. More generally, biological noise and variability have implications in a wide range of brain mechanisms such as behaviours [Faisal et al., 2008], computation [Findling and Wyart, 2021], information processing [Destexhe, 2022] or for brain dynamics [Laing and Lord, 2009]. Here, we want to model the variability of cell firing properties among a given firing type. Such inter-type variability has a biological basis, but from a modelling perspective, it originates from a trade-off one has to make to define a few representative cell types with specific firing types. Indeed, cells often form a ‘continuum of types’, as visible from the often blurry limits where some cells cannot be consistently classified [Gouwens et al., 2019, Scala et al., 2021]. In particular, firing variability has been shown to be important for a range of biological mechanisms of the brain, such as for network properties [Aradi and Soltesz, 2002], resilience to changes in synchrony with epilepsy [Santhakumar and Soltesz, 2004, Hutt et al., 2022, Rich et al., 2022], increased information content for efficient population coding [Padmanabhan and Urban, 2010, Tripathy et al., 2013, Chelaru and Dragoi, 2008, Mejias and Longtin, 2012] or energy efficiency [Deistler et al., 2022].

A cell type definition should not only account for specific values of certain electrophysiological features but also for their variability. Within a cell type, this variability will differ across species, age, or even brain regions and must therefore be factored in. In fact, for modelling studies ranging from detailed single cells to circuit simulations, such a complete characterisation of cell types will play an important role. First, for singlecell modelling, the unknown implications of the feature variability on ion channel conductances can be quantified by considering an ensemble of models. This ensemble of models will be less susceptible to containing bias than a fixed set of conductances [Marder and Goaillard, 2006]. Second, one can study the interplay between these conductances and the resulting firing types to gain a theoretical understanding or even propose some experimentally verifiable modelling hypotheses. Third, having a large population of valid, but generic models, allows the selection of sub-populations with specific properties. Constraints can be imposed directly by restricting certain feature values or indirectly via constraints on certain properties of larger circuit models, or through generalisation to other morphologies. In this study, we are interested in the latter, and in particular in defining a set of models that accurately reproduce the desired firing type on a population of morphologies taking into account its variability. This facilitates the statistical analysis of interdependencies between morphological and electrical properties. It also allows us to address the question of which morphological features affect electrophysiological properties. Finally, and most importantly, this approach allows the generation of a population of cell models with controlled morpho-electrical variability.

Several attempts to quantify the variability in neuronal parameter space required to ensure the cell remains in a specified firing type have been made in the past. It started as early as Foster et al. [1993] with systematic treatments more than a decade later. First, in Prinz et al. [2003] they build models of Lateral Pyloric neurons with this sampling approach to bypass the common hand-tuning of parameters or later in Taylor et al. [2009] to perform a more in-depth analysis of ion channels interactions in biophysical neuronal models. In these works, they randomly sample the parameter space and subsequently filtered out models with features out of their prescribed range. Other approaches leveraging optimisation algorithms such as Achard and De Schutter [2006] for Purkinje cells to understand the shape of the neuron parameter space were attempted.

Here, we will instead use the Markov chain Monte Carlo (MCMC) method (see for example [Gilks et al.,1995]) to sample the parameter space of the electrical model of rat cortical layer 5 pyramidal cells [Reva et al.,2022]. This method, already used in a similar context by [Wang et al., 2022] provides a Bayesian framework for sampling parameters of the model. It also improves on random sampling by preventing evaluations of too many models with wrong firing properties and provides statistical guarantees that the parameter space is well sampled with respect to a given probability distribution, implemented here from the cost function constructed from electrophysiological features. From these sampled models evaluated on a single reconstructed morphology, we develop a method to generalise them to a population of morphologies inspired from Hay et al. [2013], by adjusting surface areas of the axon initial segment (AIS) and soma based on relative input resistances between them and the dendrites. With these models, we studied the morpho-electrical variability of these models, specifically their ability to reproduce experimental variability.

## Results

### Experimental morpho-electrical variability

We considered two datasets: 64 morphologically detailed reconstructions of layer 5 pyramidal cells (L5PC) [Reimann et al., 2022, Markram et al., 2015] and 44 electrophysiological patch clamp recordings of the same cell type [Reva et al., 2022, Markram et al., 2015]. The L5PC morphologies are classified into four subtypes according to the properties of the apical dendrite: thick-tufted, bi-tufted, small-tufted and un-tufted (see Sec. A and [Kanari et al., 2019]). We will refer to tick-tufted the subtype denoted as TPC:A in [Kanari et al., 2019].

To illustrate the variability present in the morphologies of the thick tufted L5PC, we extracted 11 morphological features per dendritic type (see Fig. 1b) and plotted representative morphologies of the mean, large, small and exemplar cells (see Fig. 1a). The first three cells were selected based on their total surface areas, and the exemplar is the cell with the proximal dendritic surface area closest to the median profile of the population (see SI. Sec.C). We observed that morphological features such as total surface areas, total lengths or the number of bifurcations shown in Fig. 1a-b vary from three to five-fold and have various correlations (see Supp. Fig. 9). These large differences within the thick-tufted morphological type show that a single morphology cannot be a faithful representation of the entire population.

**Figure 1:**
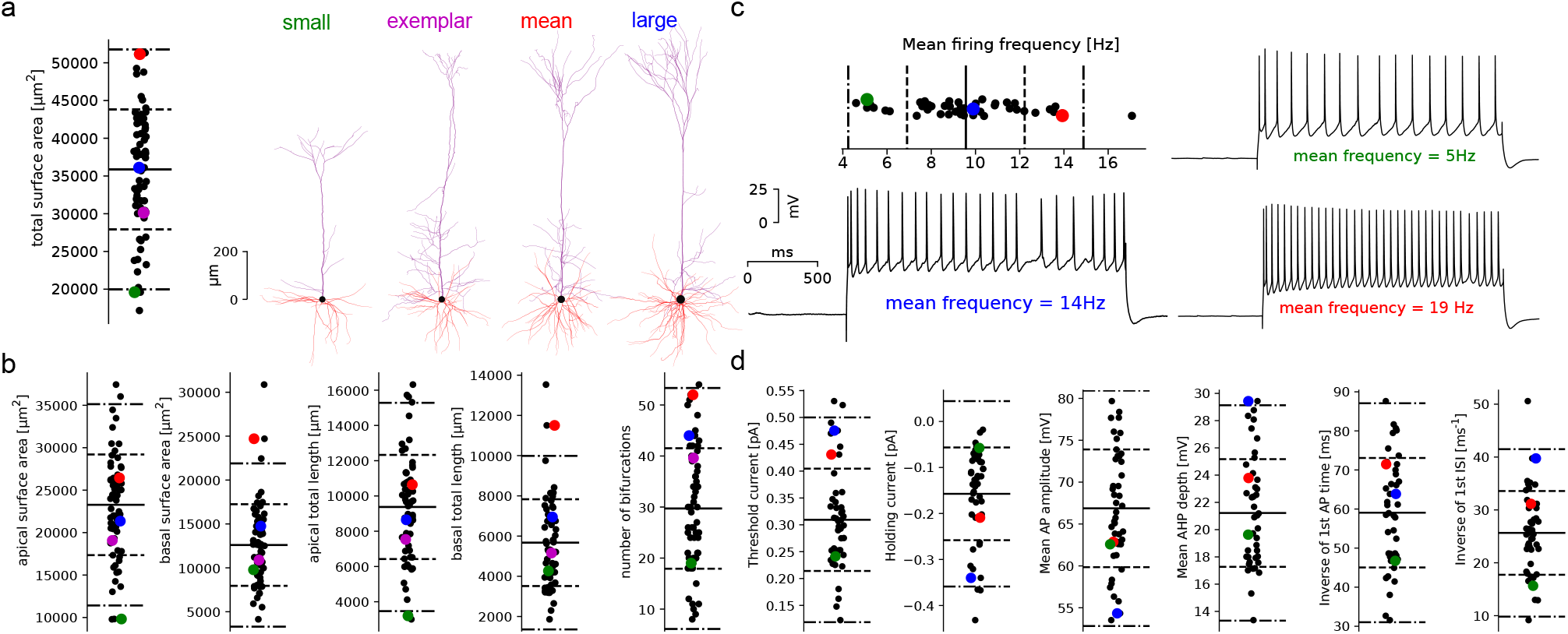
Morphological and electrical variability in thick tufted L5 pyramidal cells. **a.** Distribution of total surface areas of thick-tufted layer 5 pyramidal morphologies and four examples with small, mean and large areas as well as exemplar morphology. Coloured dots on this panel and on panel **b** correspond to these morphologies. The thick line is the mean, dash lines are 1sd and doted-dash lines are 2sd of the data (same for all panels). **b.** Distributions of some morphometrics related to morphological size and surface area. Refer to Fig. 9 for more morphometrics and correlations between them. **c.** Distribution of mean firing frequencies for the 44 recordings of step protocol at 200% threshold current, with three examples on the right, with small, mean and large mean firing frequencies. Coloured dots on this panel and on panel **d** correspond to these recordings. **d.** Distribution of some other electrical features extracted from the same protocol as well as the threshold and holding current. Refer to Fig. 8 for more features and correlations between them.

To illustrate the variability of electrophysiological properties of L5PCs we plotted the distribution of some of the 61 extracted features in Fig. 1c-d. Features are extracted from traces obtained during the experimental application of specific protocols Reva et al. [2022], Markram et al. [2015], and we chose as an illustration to use the protocol of 200% rheobase current step. In the example in Fig. 1c, we selected three recordings from three cells with different mean frequencies for this step protocol. The mean frequency range from 5Hz to 14Hz, a nearly three-fold range. The variability of other electrical features, such as holding current (required current to hold the cell at −83 mV), threshold current or other firing properties is also large (see 1d).

Overall, a few correlations are observed between morphological features (see Supp. Fig. 9, and similarly for electrical features (see Supp. Fig. 8). For example, cells with high mean firing frequency have shorter inter-spike intervals (ISI) and the cells with longer apical dendrites have larger apical surface areas. These observations suggest that, as is the case for morphological classes [Kanari et al., 2019, 2022], electrical features cannot be described purely by a set of linearly dependent features. In the case of morphologies, simple morphometrics cannot sufficiently describe the complexity of branching structures [Kanari et al., 2018] while for electrical features, the non-linear, voltage or calcium-dependent dynamics of the ionic channel conductances create a complex interplay between ionic currents Marder and Taylor [2011].

### MCMC sampling of electrical models

To reproduce the experimental variability of electrical features in our dataset we built multi-compartmental electrical models composed of the examplar morphology and a set of 30 free parameters based on Hodgkin-Huxley mechanisms, as described in [Markram et al., 2015, Reva et al., 2022] (see also Suppl. Sec. D). We then apply the Markov Chain Monte-Carlo method to sample electrical models in this parameter space as follows.

First, to assess the validity of a given set of parameters **p** to reproduce a target neuronal type, we compared the feature values extracted from simulated traces under specific protocols with the mean and standard deviation of the same features on the population of experimental recordings (see Suppl. Sec.D). More precisely, we computed an absolute z-score for each evaluated feature and a global cost function as the largest score across all features. Often, the sum of the scores is used as a cost *C*(**p**) to quantify the quality of an electrical model (see [Van Geit et al., 2016]). Here, instead, we used the maximum score as it results in overall better models by preventing any z-score from growing too large relative to the others. From the cost function, we defined a probability function on the parameter space

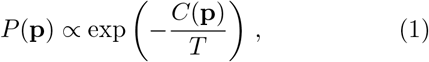

parameterised by a temperature parameter *T*. For lower *T*, the generated samples remain around local minima of the cost function while for larger *T* the samples cover more volume with larger costs. In the extreme of *T* → 0, we theoretically recover the global minimum, and *T* → ∞ leads to a uniform sampling of parameters. Sampling from our cost function ensures that most models have a low cost, with variability similar to experimental variability. To sample from this distribution, we ran several MCMC chains from random initial conditions, which we updated using the Metropolis-Hastings algorithm with multi-variate Gaussian prior (see Sec. E). The sampling quality was validated with an acceptance rate above 50% and fast decaying autocorrelation (see Suppl. Fig. 7).

Using this MCMC sampling method of the parameter space, we obtained 273’088 models, from which 209’653 have costs below our threshold of 5sd. We checked whether the obtained population of models reproduced the experimental variability of the electrical features (Fig. 1c-d). Most of the distributions are centred around the experimental mean (solid lines in Fig. 2b), except for the mean action potential (AP) amplitude, inverse time to first AP and mean afterhyperpolarization (AHP) depths (Fig. 2b). For mean action potential (AP) amplitude and inverse time to first AP, as some models are close to the mean experimental value, it could be possible to perform a specific selection for these features to obtain a distribution closer to the experimental population. However, for the mean AHP depth, all models present a smaller value than the mean experimental value, showing a limitation of our modelling approach to reproduce this specific feature.

**Figure 2:**
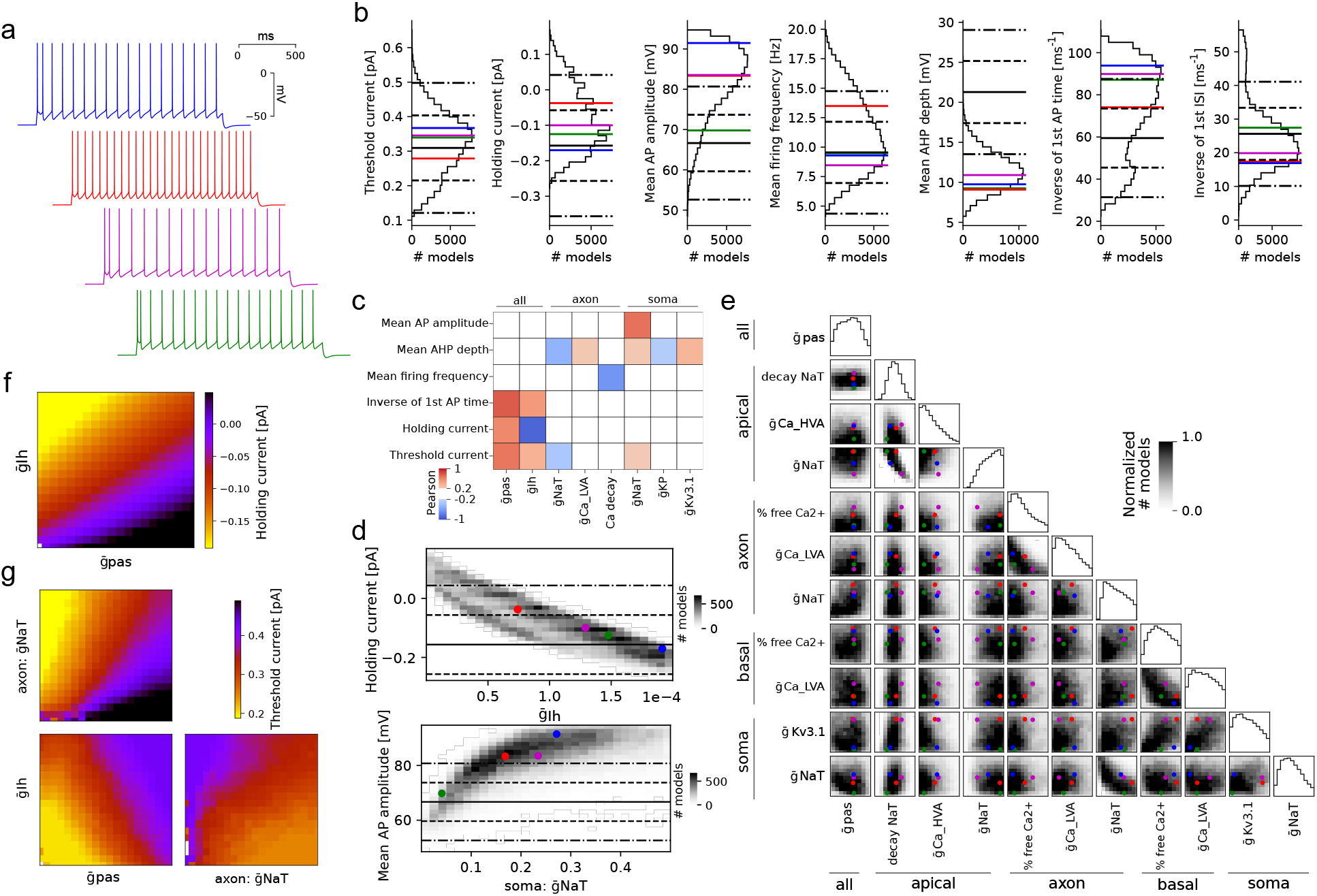
Electrical model generation with MCMC. **a.** Traces of the step 200% protocol for four models picked such that the blue has large AP amplitudes, green has small amplitudes while red and magenta have amplitudes around the median. We refer to Fig. 14 for currentscape plots of these traces. **b.** Distribution of some feature values of step 200% protocol for models sampled via MCMC with cost < 4. The thick line is the mean, dash lines are 1sd and doted-dash lines are 2sd of the experimental data (same for all panels). **c.** Correlation matrix between features of panel **b** and parameters mostly correlated with these features (Pearson >0.2). **d.** Density of models for holding current and mean AP amplitude as a function of their mostly correlated parameters, selected from **c**. The coloured dots represent the four models in **a**. **e.** Corner plot of model densities with one-dimensional marginals on the diagonal. The grey scales are normalised per pair of parameters and only the most correlated (MI>0.03) pairs of parameters are shown. The coloured dots represent the four models in **a**. **f.** Average holding current value over all parameters except for the two mostly correlated parameters to predict holding current (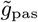 and 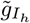). **g.** Average threshold current value over all parameters except the three mostly correlated parameters to predict threshold current (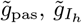 and axonal 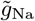).

The large population of models gave us the opportunity to explore the causal link between the model parameter values and the feature values. For this reason, we looked at the correlation between a subset of features and parameters (Fig. 2c) measured with the Pearson coefficient. We found that some features have a large correlation with few parameters (with up to 0.87 between holding current and 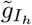 in Fig. 2d(top) or 0.68 for mean AP amplitude and somatic Na in Fig. 2d(bottom)). On the contrary, other parameters (such as AHP depth) present low correlations with a large subset of parameters (see AHP depth in Fig. 2c). Therefore, AHP depth is controlled by more parameters than other features, making it intrinsically harder to control. While, holding current is highly correlated with 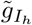 conductance,(Fig. 2d(top)), the variability of this feature cannot be fully explained by the variability in 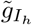. The additional variability partially comes from 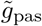, which is also correlated with holding current (Fig. 2f). In general, the variability of feature values depends on the interplay between several underlying parameters, and more parameters are required for features related to spiking (Fig. 2g and 13).

This study of the causal link between parameters and features allows a refinement of the MCMC sampling. For example, the correlation between the mean AP amplitude and the maximum conductance of the somatic sodium channel 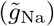 Fig. 2d(bottom) suggests a way to control the AP amplitude by adjusting the upper bound of the somatic 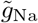 to around 0.15. In this way, most models will be more centred around the experimental mean (around 68 mV) during MCMC sampling, thus producing more valid models for the same computational cost than sub-sampling. We also looked if constraining the models below 3sd for all the features produces the emergence of correlations between parameters (Fig. 2e). For example, we observed a negative correlation between maximal somatic Na conductances and maximal axonal Na conductances (as already noticed in [Schneider et al., 2022]). This correlation is partly imposed from the constraint AP amplitude. In fact, if the somatic Na conductance is high, the axonal conductance has to be low to maintain AP amplitude within the experimental range. This representation of the accepted models (constrained by the cost) can be seen as a global map of the parameter space where valid models are located (with costs below 5sd). Regardless of their locations, the models will globally perform equally well (see Fig. 2a-b) but will contain subtle differences that can be quantified or controlled with MCMC sampling.

### Generalisation of electrical models to a population of morphologies

To sample valid models with MCMC we used a single exemplar morphology assumed to represent an entire population of morphologies. Since our original motivation was to obtain a population of models and morphologies such that any pairs were valid with high probability, we needed to ensure that our MCMC models remained valid on the entire population of morphologies. First, to check the validity of models, we used a cost function based on a reduced set of features, discarding features based on backpropagating action potentials (bAP), which are sensitive to the shape and length of the dendrite tree. Then, due to the large number of models obtained from MCMC, we began by randomly sampling 100 models with a cost below 3sd (out of 10’678), such that the MCMC density of models was preserved. This results in models being more likely to be away from the region of invalid models in the parameter space, hence possibly more generalisable. Indeed, one expects that if a model is close to the boundary of this region, a change in the morphology would affect the effective conductances of the whole cell, which may bring a cell out of the valid region.

To improve the generalisability of our electrical models on various morphologies, we adapted the soma and AIS surface area by computing the relative input resistances between soma, AIS and dendrites. Indeed, as it was noticed in early works such as [Rall, 1959], or more recently in [Hay et al., 2013, Reva et al., 2022], the relative input resistances between the AIS, soma and dendrites are essential to determine the excitability of the cell. The important quantities to consider in this context are called *ρ* factors [Rall, 1959] and are defined as the ratio of input resistances of specific compartments as

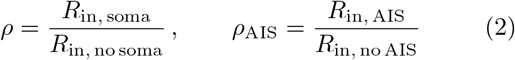

where *R*_in,soma_ and *R*_in,AIS_ are the input resistance of the isolated soma and AIS compartments, and *R*_in, no soma_ is the input resistance of all the neurites measured at the soma location (numerically evaluated at the AIS), but without the soma. Similarly, *R*_in, no AIS_ is the input resistance of the soma and dendrites, without the AIS. In [Hay et al., 2013], the authors rescaled the maximal conductances of the AIS and soma to match the *ρ* factors of the optimised cell. Here, we instead consider that the AIS and soma sizes have variability that can be exploited to improve the generalisation of fixed electrical models. We thus rescaled the surface area of the AIS and soma of each morphology such that the *ρ* factors match a target value. Our dataset of morphologies presents 4 morphological types and we extracted the examplar morphology (based on the median proximal surface area profile, see SI. Sec. C) for each of these types. We had to find the target values for *ρ* and *rho*_AIS_ for each pair between the 100 electrical models and the 4 examplar morphologies. The target *ρ* values are found with a grid search on AIS and soma scales for the optimal cost (see Sec. H). We thus have one set of the target *ρ* factors per model and per morphological type present in the population.

We first show how rescaling the AIS and soma size changes the cost of a model (Fig. 3a) on the four morphologies shown in Fig. 1. We observe that for all morphologies, a reduction of the AIS size leads to a large increase in cost (clipped at 8sd) while the change of soma size does not impact the firing pattern (Fig. 3b). The exact shape of the level set of cells with costs less than 5sd largely depends on the morphology (Fig. 3a, the region below 5sd is enclosed in the dashed lines). For some morphologies, the original AIS and soma size do not produce a valid model ((Fig. 3a, color dots). In particular, the region of valid models is small for large cells but large for small cells. By converting the AIS and soma scales to *ρ* factors (Fig. 3c) we confirm that the target *ρ* factors (black cross) obtained from the exemplar cell are within, or close to the validity region of other morphologies. If the cells are too different, such as our small and large morphologies (see Fig. 1a), their validity region in the *ρ* factor plane may not overlap for both to work with a single target *ρ* factor. As this overlap depends on the electrical model, it defines its generalisability on a population of morphology (see next section).

**Figure 3:**
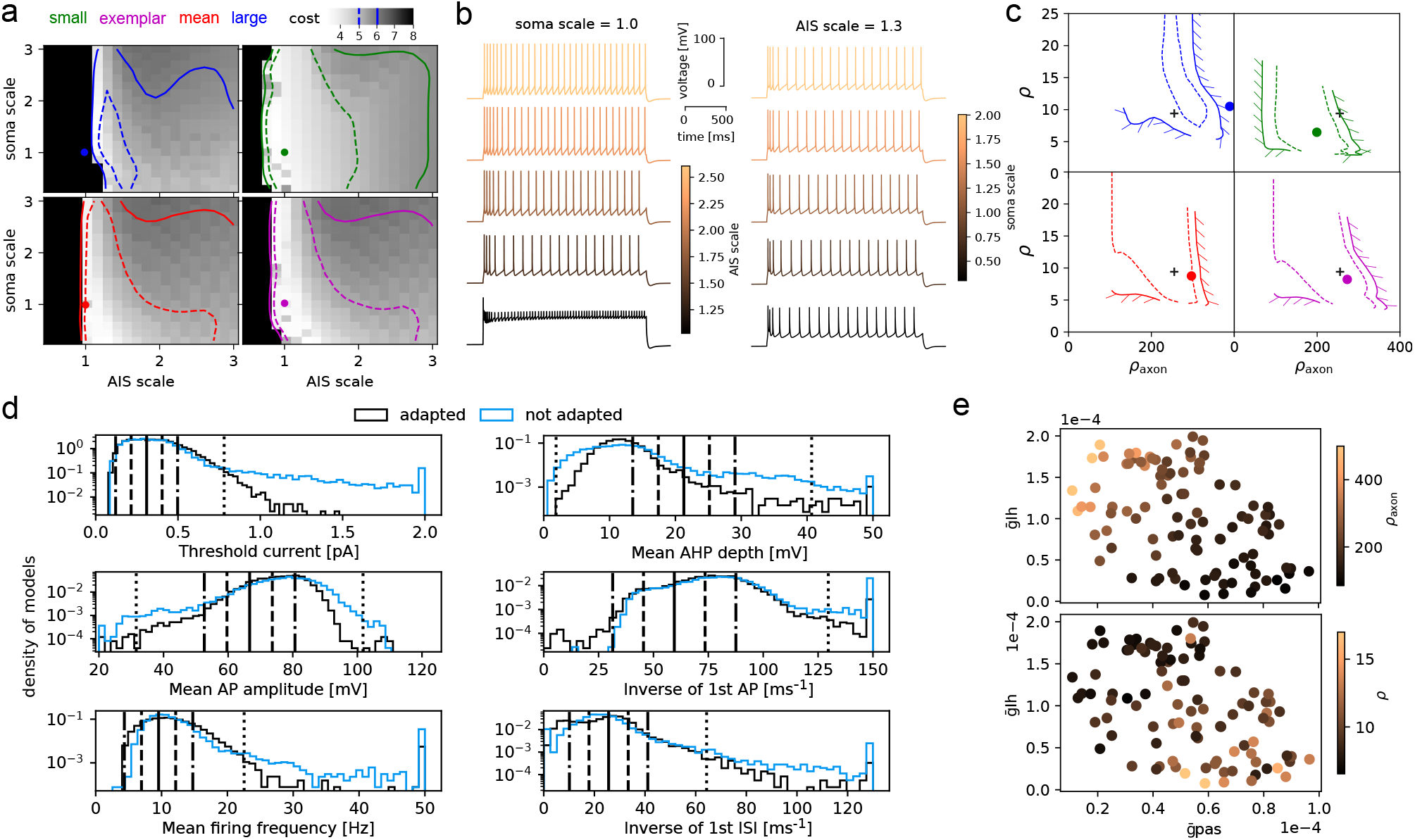
Generalisation with AIS/soma adaptation. **a.** Cost of models by varying soma and AIS size for the four morphologies of Fig. 1a. The region enclosed in dashed lines has a cost below 5sd while that in solid lines has a cost below 6sd. The dot shows the location of the cell with unscaled AIS and soma. **b.** Step traces of the large morphology with fixed soma (left) and AIS (right) scale but with varying AIS (left) and soma (right) scales. **c.** The model validity region with cost of 5sd (dashed) and 6sd (solid) of panel **a** is mapped to the *ρ* factor plane. The dots again represent the unit AIS and soma scales while the black cross is the location of the best model obtained by minimising the cost from the exemplar with the grid search in panel **a**, bottom right. **d.** Histograms of feature values on all pairs of electrical models and morphologies with (black) and without (blue) soma and AIS adaptation. The thick line is the mean, dash lines are 1sd and doted-dash lines are 2sd of the experimental data. **e.** *ρ* and *ρ*_axon_ as a function of the 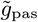 and 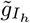 parameters.

For each model, we obtained target *ρ* factors using the examplar morphologies and we used the targets to fit the AIS and soma size of each morphology (see Sec.H). These adapted cells have a clear reduction of extreme values of features as compared to non-adapted cells (Fig. 3d), and in particular for threshold current, where 7.4% of the cells have a threshold above 5sd without adaptation while only 1.1% of cells are above 5sd after adaptation. As *ρ* factors are based on input resistance, they are highly sensitive to the passive leak and *I_h_* channels densities (Fig 3e). It should therefore be possible to predict the target *ρ* values directly by looking at these parameters instead of computing the AIS and soma scaling on the examplar morphologies. To demonstrate it, we fitted a regressor learning model (see Sec. J and H) to predict the values of *ρ* factors from the model parameters and achieve a 10-fold accuracy for the prediction of *ρ* of 1.36 ± 0.34 and for *ρ*_AIS_ of 27.5 ± 8.6. As described in the next section, this AIS/soma adjustment, which corresponds to some artificially controlled variability, is important to ensure a model generalises to a population of morphologies with substantial variabilities.

### Morpho-electrical selection and variability

Even with AIS and soma adaptation, it is not guaranteed that all pairs of models and morphologies will work together. For example, the morphology with the smallest area (blue colour Fig. 3.c) is close to the non-valid regime. It is possible that a better choice of the target *ρ* factors exists, but given our current algorithm based on a single exemplar morphology, the success rate for model morphology pairs is satisfactory, as shown in Fig. 4a. We distinguish three levels of models: valid models (light grey), models for which some features with scores larger than 5sd (grey) and models where the search for threshold current failed, hence they do not spike during step protocols (black) (Fig. 4a). By sorting the electrical models from the less generalizable (failing when applied to a lot of morphologies) to the most generalizable (produce good cells for all the morphologies), we were able to select a large fraction of morphologies and electrical models (delimited with orange lines) such that most pairs have a cost below 5sd. In Supp. Fig. 15a, we show the same selection matrix without adaptation, which needs a more drastic selection of models and morphologies.

**Figure 4:**
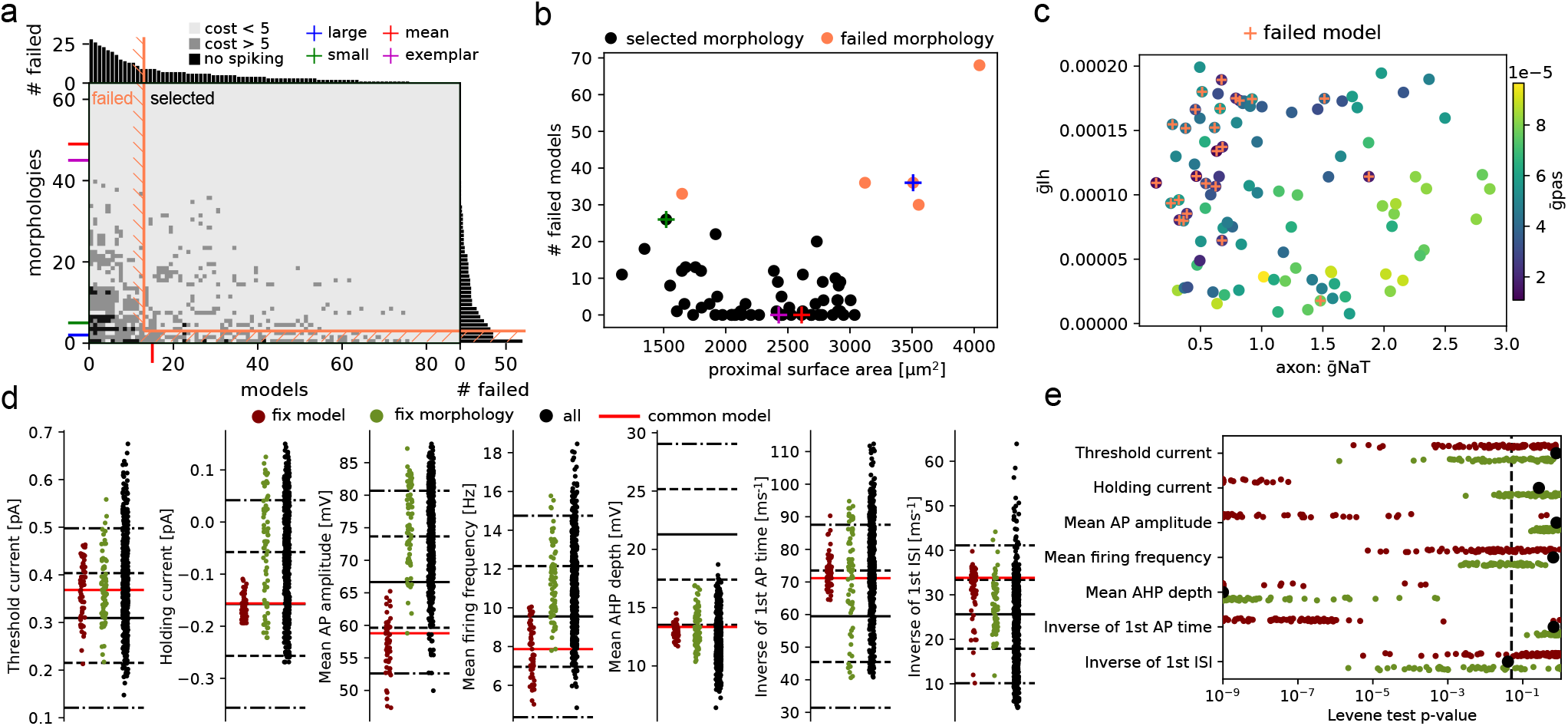
Morpho-electrical selection and variability. **a.** Selection matrix for thick-tufted cells where selected models and morphologies are delimited with the orange line. Each pixel corresponds to a model, where models in light grey have scores below 5sd, in grey have some features with costs higher than 5sd and in black pixels are models which do not spike. The four morphologies of Fig. 1 are represented with coloured ticks, and the model of Fig. 4 with a black tick. We refer to Supp. Fig. 15 for the same selection matrix without AIS and soma adaptation. **b.** Proximal (up to 500μm) dendritic surface areas for all morphologies with non-selected morphologies in orange versus the number of failed models for each morphology. The four morphologies are also highlighted with coloured crosses. **c.** From the classification of selected models from their parameters with a classifier algorithm (10-fold accuracy of 0.89 ± 0.08), three parameters are most important, with a clear correlation. Models with small 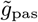 values also contain models for which we have not been able to fit the AIS/soma resistance model (not shown in **a**). **d.** Distribution of features obtained by freezing the black model (57 points in red), the exemplar morphology (75 points in green) or 500 models randomly sampled from the 5197 total pairs with cost < 5 (in black). The red line corresponds to the value of the black model with exemplar morphology. The thick line is the mean, dash lines are 1sd and doted-dash lines are 2sd of the experimental data. **e.** p-value of Levene’s test centred with mean. Each dot is the p-value of this test between the experimental data shown in Fig. 1 and all MCMC models with cost < 3 in black, subsets of models in red and morphologies in green. The distributions per feature in panel **d** corresponds to single points in this plot.

Given the morphological variability of our population of pyramidal cells, we can investigate the relationships between some global morphometrics and generalisable cell models. For example, the proximal (up to 500μm) surface areas of selected (black) and nonselected (orange) morphologies is a good predictor of the number of failed models, of model generalizability (see Fig. 4b). In fact, best morphologies (i.e that failed with a small number of electrical models) have surface areas close to the exemplar morphology, while most distant morphologies (with large or small surface areas) are much less generalisable (they failed with a large number of electrical models). This result indicates the range of proximal surface areas of morphologies that can be generalisable from a single exemplar morphology. We also show the same result without adaptation where the region of validity is below the exemplar ( Suppl. Fig. 15b). If the largest cells are important for a given study, two choices are possible. One can recalibrate the *ρ* factors using a larger exemplar morphology, or, if this is not sufficient, run MCMC sampling again to produce another set of models. The second choice will arise only for extreme morphological differences where the valid regime in the parameter space is not reached with the MCMC sampling based on the first exemplar.

In this context, we ask whether it is possible to predict if an electrical model is generalisable from the parameter values. From inspecting parameter distribution between each set of models (generalisable and not generalisable) there are no evident differences (not shown), but when training a machine learning classifier (see Sec. J), the 10-fold accuracy reaches 0.89 ± 0.08, showing that it is possible to accurately predict the generalisability of a model on a given population, but only via nonlinear, higher-order combinations of parameters. In Fig. 4c, we show the values of the main parameters involved in predicting the generalisability of models on the population of morphologies. We found that small passive and axonal Na conductance and large *I_h_* conductances are more likely to produce non-generalisable models.

Finally, from this population of models and morphologies, we assessed how well they match the variability of the experimental data (Fig. 1), and in particular, the morphological or electrophysiological variability alone is sufficient to reproduce the experimental variability. For this, we fixed a model and compared the distributions of the features when evaluated on all selected morphologies (Fig. 4d), red dots). We found that, for this specific electrical model, the morphological variability is not sufficient to explain the experimental features variability. In fact, we can see that for features such as holding current, AP amplitude or time to first AP, distributions of features obtained by modification of morphology are less variable than experimental features (Fig. 4d, the thick line is mean, the dashed line is 1 sd and the dotted dashed line is 2sd of experimental data). Then, we fixed a morphology and compared the distributions of the features when evaluated on all selected electrical models (Fig. 4d, green dots). Fixing a morphology seems to produce more consistent variabilities across features, except for AHP depth, which is biased towards low values already in the MCMC sampling. Therefore, the experimental variability of electrical features seems to mostly arise from the variability of the ion channel densities in the population. We finally looked at the feature distributions when testing all the pairs between selected morphologies and selected electrical models ((Fig. 4d), black dots) and found an even larger feature variability, showing that combining both morphological and electrophysiological variability is important to reproduce experimental variability. In order to quantify these variabilities, we compared these distributions with the experimental data with the Levene statistical test (centred with mean) (Fig. 4e) by fixing all morphologies (green) or all models (red). We found with this procedure, more feature distributions presented a p-value for the Levene-test smaller than 0.05 with a fixed electrical model than with a fixed morphology, consistent with the results in panel d. Finally, the black dots show the comparison between experimental feature distributions and distributions when testing all the 5197 pairs between electrical models and morphologies which all have p-values above 0.05 (distributions not distinguishable from the experimental one) but for the mean AHP depth. Therefore, applying MCMC sampling, soma and AIS scaling by *ρ* factors procedure and selection of generalizable electrical models and morphologies, allowed us to build a population of models that reproduces the variability of the features found in a neuronal population.

### Further generalisations

From MCMC sampling, we only selected 100 models to perform AIS/soma adaptation to create a set of valid pairs of electrical models and reconstructions. This was primarily due to the computational cost of calibrating the *ρ* factors and evaluating all the models on morphologies to select them. By using standard tree-based machine learning regressors and classifiers (see J) we fitted models for the *ρ* factors, the AIS/soma resistances and model generalizability, we were able to use more models from MCMC sampling with reduced computational cost. In total, we evaluated 13794 new pairs of selected morphologies and sampled models where 92.8% have cost below 5sd (as light grey pixels in Fig. 4a) and 98.9% that are able to fire (as grey pixels in Fig. 4a). Once calibrated, our method of adapting the soma and AIS can be applied to models sampled with MCMC not yet used and produce a larger number of generalisable models without the need for expensive calibration and validation of *ρ* factors and fully leverage the variability created by MCMC sampling for statistical analysis or circuit building.

Increasing the number of models is often not sufficient to capture the entire biological variability, especially if experiments involve inter-neuron connectivity or synaptic inputs. For this, one could use more reconstructions if they fall within the estimated valid surface area bounds, or leverage neuronal synthesis algorithms. Here, we generated morphologies with the algorithm of [Kanari et al., 2022] based on the topological descriptor introduced in [Kanari et al., 2018] (see SI. Sec. K) to show that if morphologies fall within the original population, there is a high probability that they will perform well on all selected models. We found that 94.8% of the generated pairs from 100 synthesised morphologies had a cost below 5sd, but only a few with small oblique surface areas were consistently failing. This suggests that some specific morphological features should also be taken into account to refine how morphologies are classified to produce consistent populations for electrical modelling.

In this work, we focused exclusively on layer 5 pyramidal cAD cells, but in Supp. Fig. 12 and SI Sec. L, we show the application of MCMC and generalisation on another electrical type, the continuous nonaccommodating interneuronal types. We found similar properties to the cADpyr models, whereby MCMC captures non-trivial parameter correlations, small morphologies are harder to generalise but from the selected set of morphologies and models, we are able to generate more valid models. Our approach of MCMC sampling and further model and morphology selection thus works on any electrical types, provided sufficient features are available and can be used to construct models with controlled variability for several electrical types, for intertype comparisons or building more realistic microcircuits.

## Discussion

In this work, we present a framework to study detailed biophysical models of neurons. In particular, we take into account the experimental variability of morphologies and electrophysiological features within cell types. For this, we leveraged a standard statistical method, MCMC, to generate thousands of electrical models reproducing experimental variability, and we generalised them to a population of detailed reconstructions of morphologies by adjusting AIS and soma size according to calibrated *ρ* factors.

With this approach, we produce a population of models with feature variability close to experimental data and we demonstrate that morphological variability alone is not sufficient to reproduce the observed electrical behaviour. The variability of electrical models as measured by the feature values is a direct consequence of the parameter variability [Marder and Goaillard, 2006]. In order to ensure that the feature variability matches the experimental data, strong constraints are necessary on the model parameter space, which is possible to be analysed with MCMC samples. The study of these constraints is out of the scope of this study but preliminary analysis indicates that a few constraints are low-dimensional, while many are high-dimensional, in particular for complex features such as average after-hyperpolarisation depth during a step protocol. These constraints on the parameter space are a result of the choice of protocols and features. Thus to obtain more specific models, reproducing more specific firing, such as BAC firing [Larkum et al., 1999, Hay et al., 2011], even stronger constraints are most certainly required such as more specific apical ion channels. Hence, such MCMC sampling methods, or improvements of them [Wang et al., 2022], provide a tool to detect model limitations and investigate their origins.

The study of model generalisability on a population of morphologies suggests a new morphological grouping based on specific morphological features, related to their electrical activity. Given an exemplar with calibrated *ρ* factors, we find that the proximal dendritic surface area (i.e. up to around 500*μm* including basal and oblique dendrites) is a good predictor for the validity of the morphology on that model. However, other more specific morphological features, such as oblique dendrite areas, also have an impact, as was discovered with synthesised morphologies in Sec. K. As such morphological features do not necessarily correlate with more generic classifications involving specific apical features (tuft, no-tuft) or axonal ones (mostly for inter-neurons), it may be possible to create more targeted morphological types of neurons based on these to ensure that all morphologies of a specific type will perform well with a single set of *ρ* factors.

Overall, this work proposes a new perspective on building detailed electrical models of neurons with Bayesian statistics via the MCMC method. We show that we can unravel subtle mechanisms via the study of parameter and electrical feature correlations while providing a consistent framework to assess the quality of electrical models by controlling their variability on a single morphology or for a population with its own morphological variability. This work opens further research avenues to study the interplay between electrical models and morphologies, co-regulation [Yang et al., 2022], energy efficiency [Bast and Oberlaender, 2021, Jedlicka et al., 2022], the impact of morpho-electro variability in circuit simulations ( [Prinz et al., 2004, Marder and Goaillard, 2006, Goaillard et al., 2009]) or links with gene expressions Bernaerts et al. [2023]. In addition, MCMC sampling of electrical models may become instrumental in making progress on the longstanding question of redundancy and synergy in biology, and in particular, in neurons [Marder and Taylor, 2011, Marder, 2011].

## Contributions

*Alexis Arnaudon:* Conceptualization, Methodology, Software, Validation, Formal analysis, Writing-Original Draft, Writing-Review & Editing, Supervision. *Maria Reva:* Conceptualization, Methodology, Formal analysis, Writing-Review & Editing, Supervision. *Michael Zbili:* Conceptualization, Methodology, Writing-Review & Editing, Supervision. *Werner Van Geit:* Supervision, Writing-Review & Editing. *Lida Kanari*: Supervision, Writing-Review & Editing. *Henry Markram*: Supervision, Funding acquisition.

## Acknowledgement

We acknowledge Karin Holm for her careful proofreading of this manuscript and Tanguy Damart for support with electrical modelling software. This study was supported by funding to the Blue Brain Project, a research centre of the École polytechnique fédérale de Lausanne (EPFL), from the Swiss government’s ETH Board of the Swiss Federal Institutes of Technology.

## Supplementary information

### A Morphology dataset

We use the layer 5 pyramidal morphology dataset of [Reimann et al., 2022, Reva et al., 2022], which is an extended version of the one from [Markram et al., 2015]. It consists of four subtypes, with 64 thick tufted cells, 38 bi-tufted cells, 30 small tufted cells and 27 un-tufted cells as defined in [Kanari et al., 2019].

### B Diametrization algorithm

The input to the algorithm is a population of morphologies, and it will be applied to a single type of neurite (basal and apical). For each section of morphology, we compute the path distance from its end to the downstream terminal point that is furthest away (see Fig. 6a. We then normalised these values by the largest path length of the morphology. We also record the section mean diameters and fit a polynomial function of diameter as a function of the distance, which will be our diameter model, see Fig. 5e.

**Figure 5:**
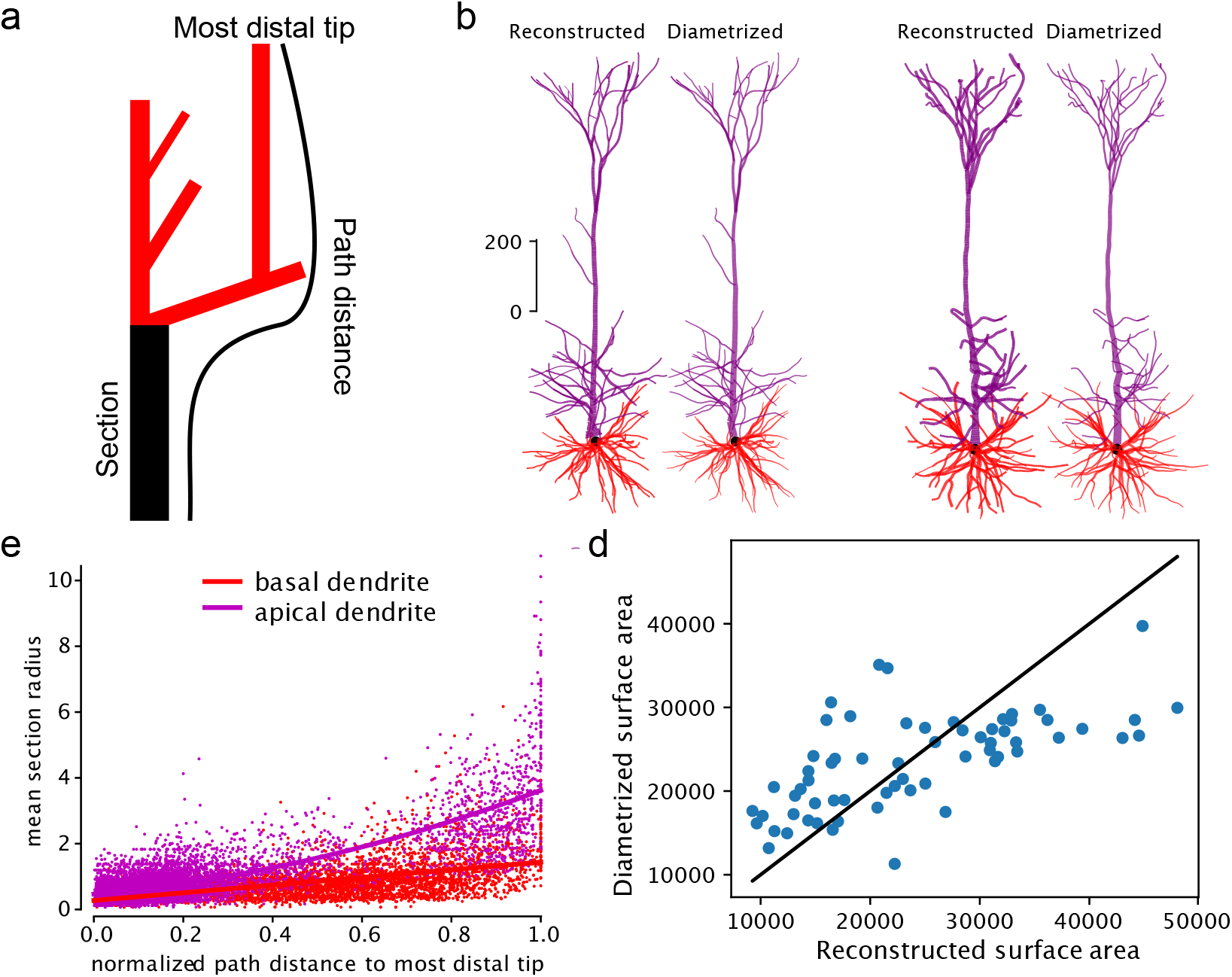
Diametrization algorithm. **a.** Illustration of distance computation used in the diametrization model for the section in black. **b.** Example of two reconstructed morphology (left) and its rediametrized versions (right) **c.** Mean section diameters as a function of path distance to the most distal terminal section for the population of thick-tufted morphologies, with a fit per type of neurite. **d.** Comparison of surface areas of all dendrites form 0 to 500*μm* between original and diametrized morphologies.

**Figure 6:**
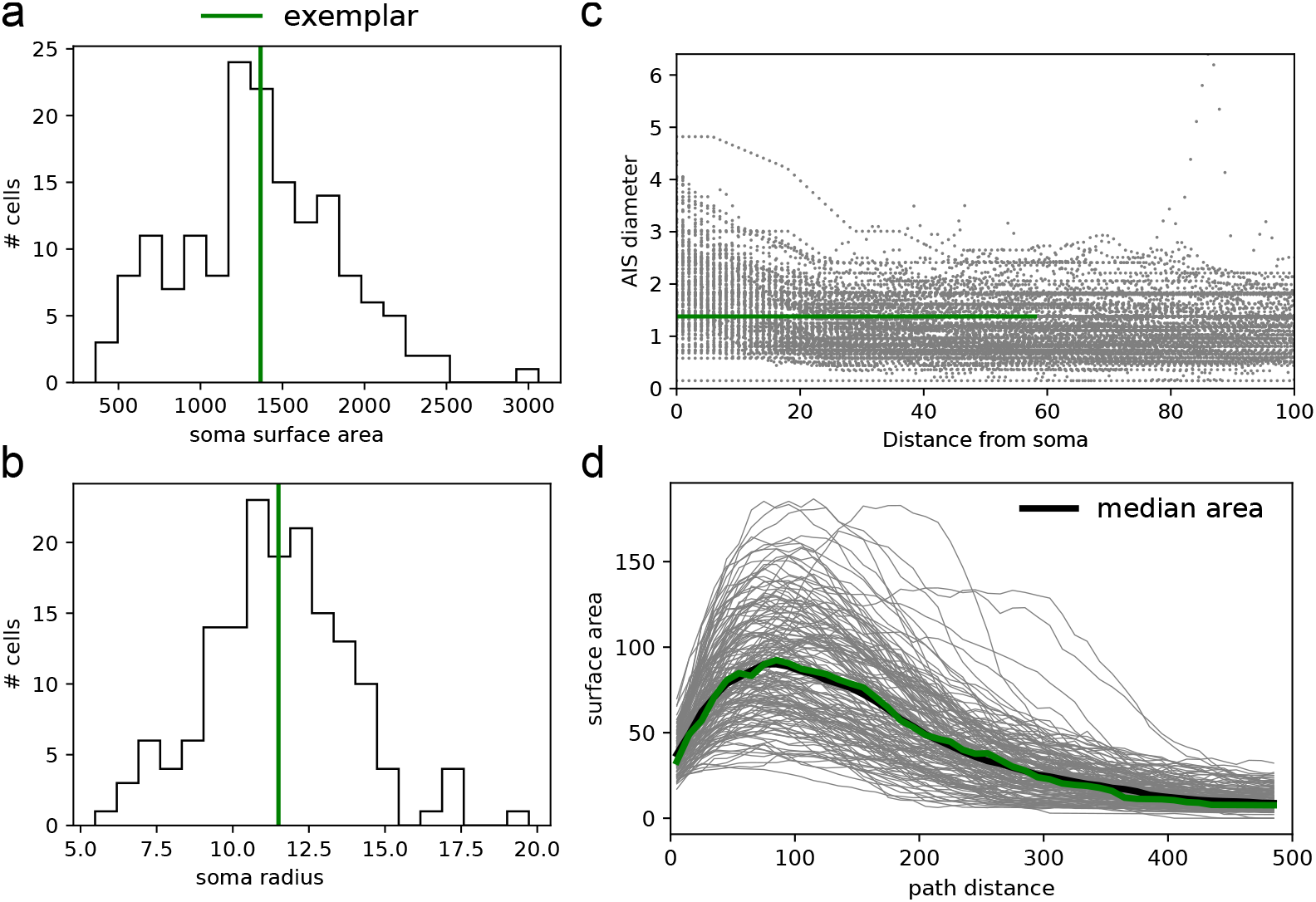
Construction of exemplar morphology. **a.** Distribution of soma surface areas of all L5 pyramidal cells, with chosen area in green. **b.** Distribution of soma radii of all cells with chosen radius in green. **c.** Diameters of the reconstructed points of the first sections of the axon, interpreted as AIS diameters. In green is the average AIS diameter of the first 60*μm* used for the exemplar. **d.** Proximal surface area profile of all dendrites, computed with 50 bins in path distance. The median profile is in black, and the closest to it is the choice of exemplar dendrites (in green).

The diametrization of a given morphology then first uses this model to assign a diameter to each section independently. It then introduces a linear tapering of diameters along each section such that the first diameter is the assigned one, and the last is the average between the first of the current section and the largest first diameter of child sections. If the section is a terminal section, we read the last diameter from the model.

In Fig. 5b we show two examples of rediametrized morphologies, where the left morphology has similar diameters, and the right one has reduced diameters. In Fig. 5d we show the changes of surface areas induced by the rediametrization, mostly affecting the large surface areas (such as for the right morphology), but overall making the distribution of dendritic surface more narrow.

The diametrization algorithm is implemented in the open-source python package https://github.com/BlueBrain/diameter-synthesis.

### C Exemplar morphology

From a population of morphologies, we select an exemplar morphology as follows.

We compute the average surface area of the soma, computed by neuron for greater consistency with Neurolucia format in Fig. 6a as well as the average radius in Fig. 6b. We then create a single cylindrical compartment with an average radius and length computed such that the surface area is also the average from the population.

We extract the diameters of the first 60*μm* of the reconstructed axons (see Fig. 6c), assumed to be the AIS of the neuron and use the average diameter to create a two-compartments model of the AIS with constant diameter. We ignore the tapering near the soma, which does not affect significantly the electrical features (not shown). We then discard the reconstructed axons as we do not electrically model them in detail but replace them with a constant diameter AIS of length 60*μm* followed by a 1 mm myelinated section. The AIS will serve to generate action potentials, and the myelinated section will act as a sink, as an approximation of the effect of removed axonal branches.

In addition, we need to select a morphology that is most representative of the population. As most experimental protocols and recordings we will use are somatic, electrical models will be most sensitive to the proximal surface area of dendrites. We compute the total surface area of all dendrites as a function of path distances in Fig. 6d, compute the median profile and select the morphology closest to use as exemplar dendrites. We apply this procedure to construct a global exemplar morphology from all L5 PC cells which we will use for MCMC sampling. We also create m-type specific exemplars where the dendrites are selected using only morphologies within this m-type, for *ρ* factor calibrations.

### D Electrical models and cost function

We use the model of [Reva et al., 2022], based on the ones from [Markram et al., 2015]. They are composed of a set of coupled nonlinear equations assigned to each compartment in a morphology. For each compartment, a subset of these equations is parametrized by properties such as radius and lengths and coupled to the adjacent compartment via current conservation condition [Hines and Carnevale, 1997]. All dynamical equations are of the form

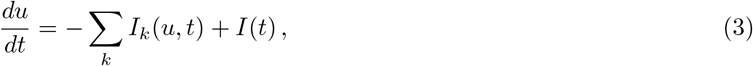

where *u* is the membrane potential, *I*(*t*) the applied current and *I_k_* the various ionic currents indexed by *k*, non-linearly depending on the voltage.

Several protocols defining *I*(*t*) are applied on the soma and recorded in the soma or along specific dendrites. These protocols are the same as in [Reva et al., 2022], where the most important is the step protocols with amplitude relative to the rheobase of the cell. The rheobase is found with a bisection search, where the lower bound is the holding current, defined as the current to hold a cell at −83mV and the current to be at −30*m*, estimated from the input resistance. The holding current is also found from a bisection search, which, as for the threshold, is terminated once a certain accuracy on the current is reached (difference between the last upper and lower bound). If the threshold current is above −30*mV*, we assume that the cell cannot spike, thus subsequent protocols are not evaluated, and the cost of the model is maximal. From these voltages (or other ion channels) recordings, a number of features, such as mean AP amplitude and AP frequency are extracted, as in [Reva et al., 2022].

Only a subset of all features is used for analysis steps after performing MCMC. In particular, we discard bAP features, which are too sensitive to apical diameters, and APWaveform where the second AP may not happen due to too short step protocol, SpikeRec because it is related to recovery after a spike, which we do not consider here and IV.

Denoting the simulated *k* feature values as a vector **f** = (*f*_0_,…, *f_k_*), the parameters as **p** = (*p*_0_,…,*p_n_*) as and construct a cost function to measure the model quality as

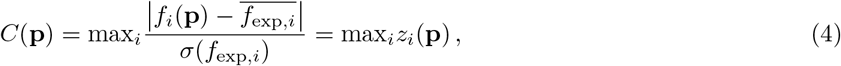

where *z_i_* are the absolute *z*-scores of each feature indexed by *i*. Notice that we define the cost as the max of the scores, which is stronger than the sum of the score usually used in optimisation [Van Geit et al., 2016]. In addition, each parameter is assigned a predefined range of possible values, possibly consistent with biological data if any are available.

### E MCMC sampling of models

All parameters have normalised versions denoted by 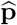, such that the available range is [−1, 1], and no bias is induced by the various possible units or bound sizes of each parameter. The parameter space for a valid electrical model is thus a subspace Ω of the hypercube [−1, 1]^*n*^ defined as *p* ∈ Ω if and only if *C*(**p**) < *C*\*, where *C*\* is a maximum cost to consider a model valid.

To generate many set of parameters to cover this set Ω, we use the MCMC method with the Metropolis-Hastings algorithm as follow. We define a probability distribution from the cost function as

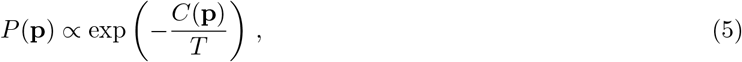

where *T* is a temperature parameter. The goal of MCMC is to sample from this distribution. For this, we use a normal prior distribution 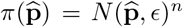 with variance *ϵ*. Because the parameter space is a hypercube, if the proposed set of parameters lies outside, we re-sample until we get a point inside the hypercube. For each chain, we then sample a random point 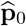 in the hypercube, and compute the next point in the chain using the Metropolis-Hastings algorithm:

1. propose 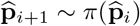,
2. draw *a* ~ *U*(0, 1),
3. accept 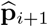 if 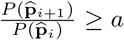, else reject and set 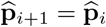,
4. repeat until number of iterations is attained.

In practice, we launch several chains for a short so-called burn-in phase so that enough chains have reached the valid region Ω. We then restart new chains from a selection of best models obtained in the burn-in phase, see Fig. 7a. To obtain our final set of parameters in Ω, we remove samples with *C* > *C*\*. The parameter ε should be set such that the acceptance rate is around 60 – 80% depending on the chain, to ensure we optimally explore *P*(**p**) correctly. The convergence plot in Fig. 7a shows the burn-in phase and the longer run, with a uniform sampling of cost values across iterations. In addition, the auto-correlation plot for this run in Fig. 7b shows fast decay of correlations after less than 50 iterations, which gives an indication of the minimum number of steps one should do to have a good sampling In Fig. 7c, we show for each feature the fraction of time it is the largest, so it defines the cost. A more uniform distribution shows that no features are primarily blocking the chains to reach low-cost values.

**Figure 7:**
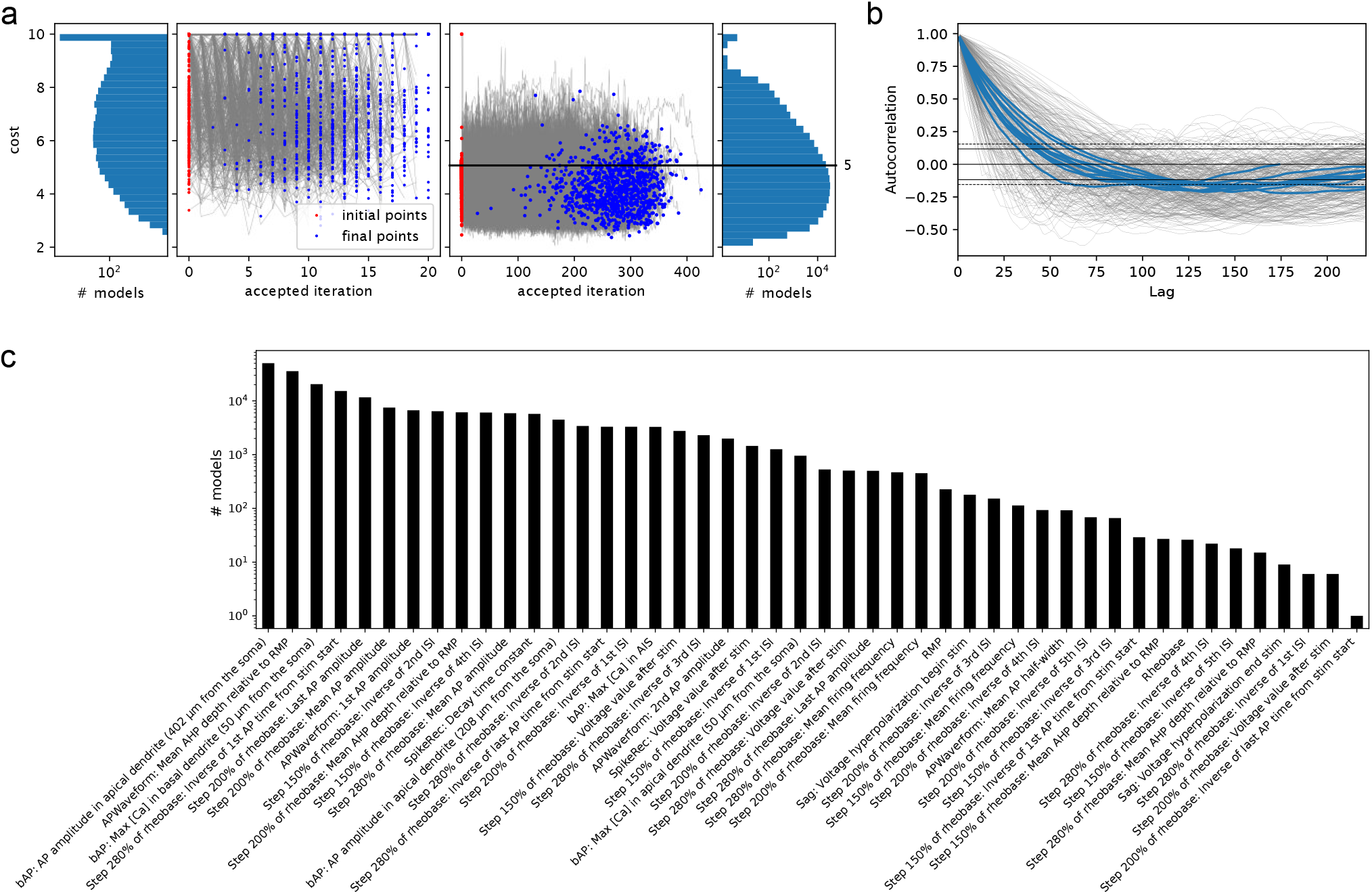
MCMC additional figures. **a.** Cost convergence for burn-in and main MCMC run. Red dots correspond to the initial costs of each chain, grey lines are the chain costs trajectories, and blue dots correspond to the final cost when the chain was stopped after 500 iterations. The x-axis corresponds to the accepted iterations, thus the horizontal scatter of blue points represents the variability in acceptance rates and an average acceptance rate of around 60%. The left and right panels are respectively the distributions of costs for the burn-in phase (left middle panel) and main MCMC run (right middle panel). **b.** Auto-correlation of all the chains in grey, and 10 chains in blue. Horizontal lines show significance intervals, as defined in Panda’s auto-correlation plot function. **c.** Distributions of scores saturating the cost for all models with costs < 5 (marked with a horizontal black line in panel **a**).

Overall, MCMC sampling can also be used as a tool to improve electrical models. Indeed, the corner plots and one-dimensional marginals such as shown in Fig. 2e is useful to adjust parameter bounds. If many good models are near the upper bound of a parameter, one could increase it (if the value remains meaningful) to maybe obtain even better models. On the contrary, if the distribution is very narrow towards small values, reducing the bounds will improve the acceptance rate of the MCMC sampling. The study of correlations between features (see Sec. F) may also be of interest to detecting the possible lack of feature parametrisability of the model, resulting in a limitation of the lowest possible scores achievable. With the addition of plots such as Fig. 7c which indicates which features saturate the cost more often, or even with plots of traces for some specific models, MCMC sampling provides can be effectively used to propose improvements on the choice of ion channel mechanisms or associated parameter bounds.

### F Correlations of features

In Fig. 8a, we compare the feature correlation computed with mutual information between experimental data (see Fig. 1) and MCMC sampling (see Fig. 2)l showing a good agreement. For example, the inter-spike interval correlation is stronger in MCMC than in the data, possibly due to experimental noise not present in our simulations. The correlation between mean frequencies is also higher in MCMC sampling, thus IF curves will have more consistent slopes among MCMC models than experimental data. In Fig. 8b we show some of these correlations via scatter plots of experimental and numerical data. In these plots, some experimental outliers (crosses) were detected and removed to compute the mutual information, as they biased the results substantially. It may therefore important to perform such analysis of the experimental data to detect any possible outliers which may not be visible in one-dimensional distributions. In addition, these outliers are few but correspond to stuck cells, thus having stuck cells in the model may be allowed in small numbers, from a pure data point of view.

**Figure 8:**
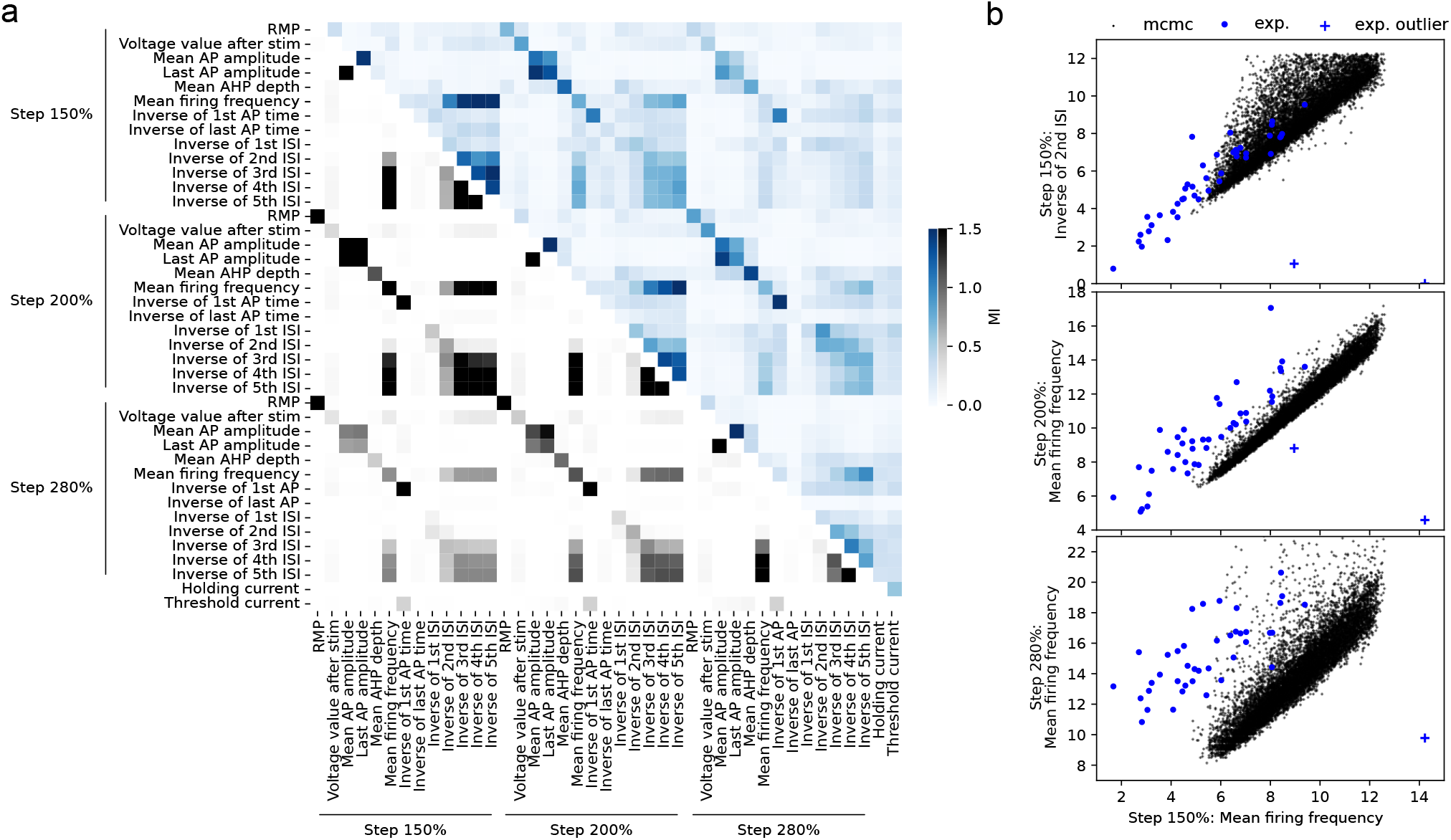
Correlations of experimental and MCMC features. **a.** Correlations (Mutual information) between features in experimental data (blue) and MCMC samples (black) for all samples with costs < 3. **b.** Scatter plots between some pairs of features, showing differences and similarities between experimental data and the MCMC sampling. Outliers have been discarded to make a correlation matrix.

In Fig. 9, we perform a similar analysis but on morphological features of reconstructed and synthesised morphologies (see Sec. K). The pairwise correlations of the selected feature show a good agreement between both sets of morphologies, where only a few features are strongly correlated.

**Figure 9:**
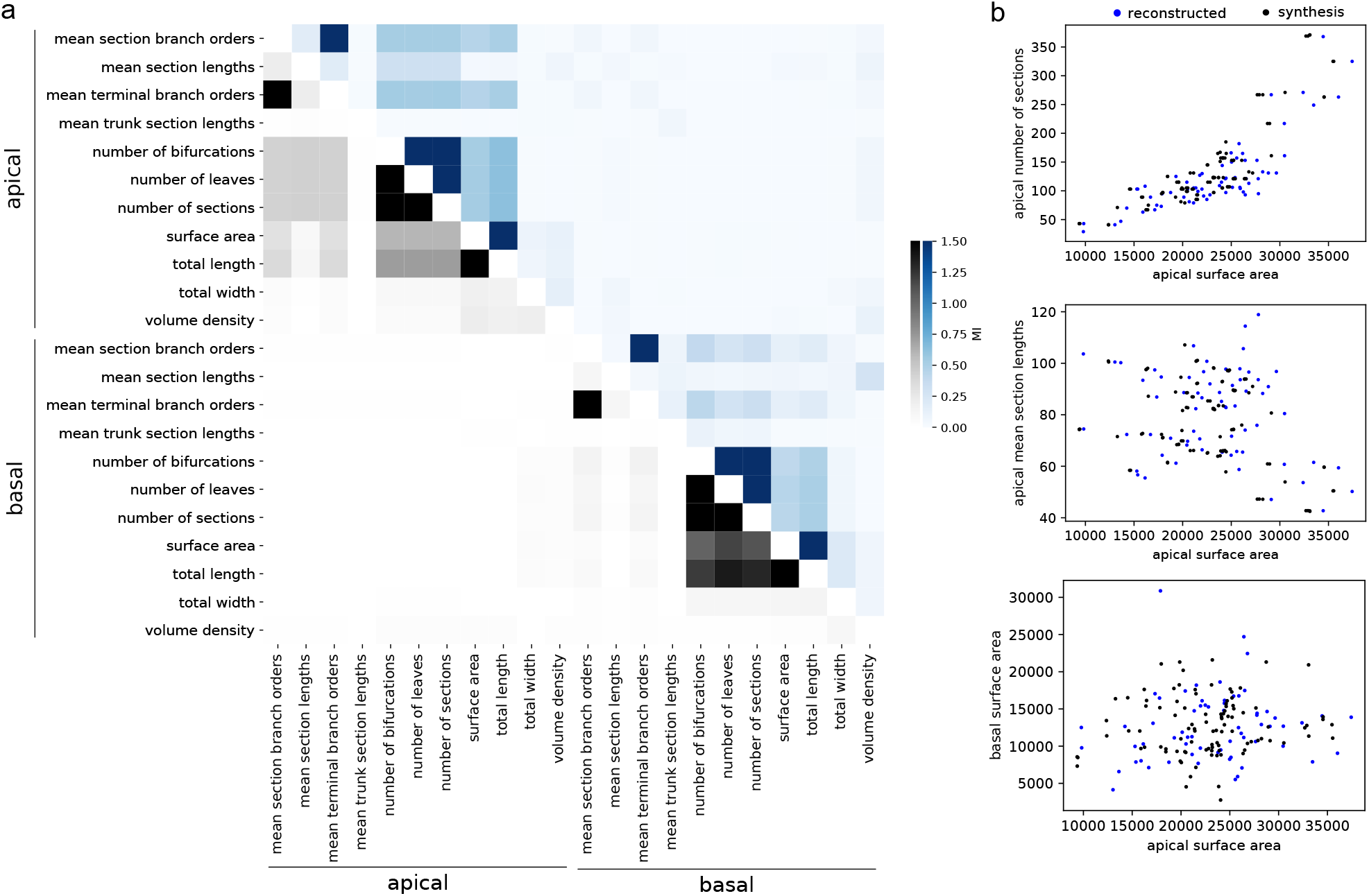
Correlations of morphological features. **a.** Correlations (Mutual information) between some morphological features in reconstructed (blue) and synthesised (black) morphologies. **b.** Scatter plots between some pairs of features.

### G Soma and AIS input resistance models

To adapt the size of soma and AIS according to the *ρ* factors, which are based on input resistances, we need to model the input resistance of isolated soma and AIS as a function of their sizes. For each model, we evaluate the input resistances as a function of the soma and AIS scales from 0.1 to 10 and perform a polynomial fit of order 3 on this data in log-log see Fig. 10a,b, resulting in four parameters per electrical model for both the AIS and soma. In Fig. 10c-h we show the parameters mostly correlated with the first three fit parameters (the fourth is also correlated with 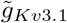), found by searching for model parameters with Pearson above 0.7 with the fit parameters. We remark that the passive currents control the affine part of the input resistances of these compartments, while the Kv3.1 current controls its deviation slope and possible deviations from a linear relation.

**Figure 10:**
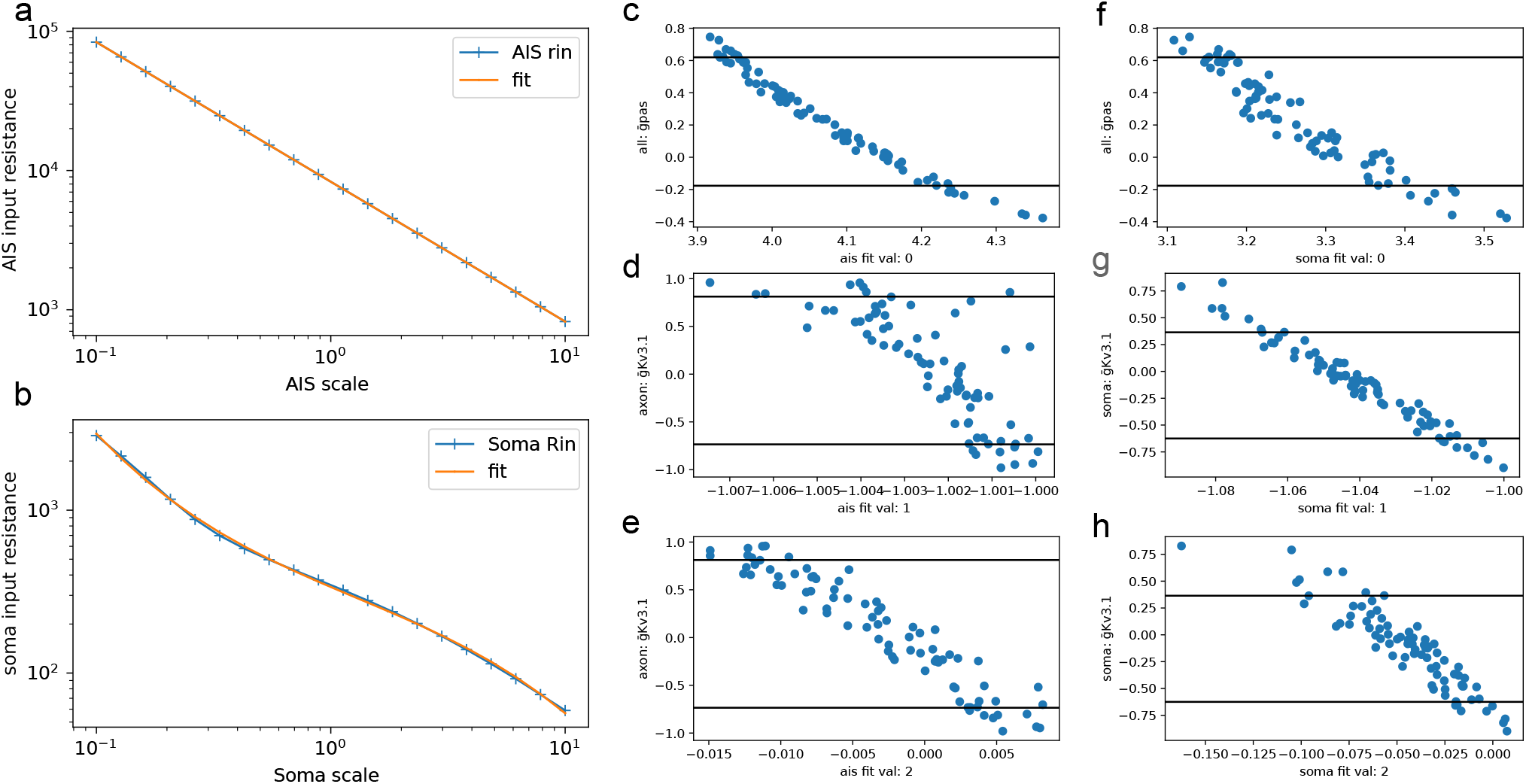
Soma and AIS input resistance models. **a** AIS input resistances for model (blue) used in Fig. 3 and a cubic polynomial fit (orange). **b** Somatic input resistances for model (blue) used in Fig. 3 and a cubic polynomial fit (orange). **c-e** Mostly correlated model (normalized from −1 to 1) parameters with the first three fit parameters for the AIS resistance model. Black lines are 10 and 90 percentile on the parameter values. **f-h** Mostly correlated model parameters with the first three fit parameters for the soma resistance model. Black lines are 10 and 90 percentile on the parameter values.

Sometimes, the computation of these input resistance models fails, either due to simulation issues, or poor polynomial fits. In this case, we discard the electrical model for later use. These cases happened more frequently for low models with low passive conductances, where other channels have more prominent effects, making the input resistance relation with the AIS and soma size more complex.

### H M-type specific rho factors

To find optimal *ρ* factors we use the exemplar morphology for each m-type and scan for AIS and soma scales from 0.5 to 1.5 in 10 steps and smooth the scores (with reduced feature set) with a Gaussian kernel of 0.1 width, to obtain a matrix such as in Fig. 3, but with lower resolution for computational efficiency. The point of the lowest score is selected as the target scale for each pair of m-type/electrical models (as the black cross in Fig. 3c), which are then converted to *ρ* factors and stored to later adapt the AIS/soma.

### I AIS/soma adaptation algorithm

To adapt the AIS and soma size to match the target *ρ* factors estimated in Sec. H, we use an iterative algorithm as follows. First, we adapt the AIS size after evaluating the input resistance of the dendrites and soma and using the polynomial fit of the AIS resistance model to assign an AIS scale. We then do the same for the soma, with input resistance computed with dendrites and scaled AIS. We repeat this two times to ensure that the scales of AIS and soma have converged. We do not require a precise convergence to the target rho factor because it has been calibrated with a small resolution for computational efficiency (see Sec. H) and nearby values are also likely to work equally well. Nevertheless, this two-step algorithm converges to *ρ* factors values at around a few per cent of the target.

### J Further generalisation with machine learning classifiers and regressors

Due to the computational cost of searching for target *ρ* factors for each electrical model and fitting input resistance models of the AIS and soma, we cannot use too many sampled models from MCMC. Hence, to further expand the pool of models, we leverage machine learning algorithms to estimate these values.

For that, we use gradient boosting tree-based learning algorithm xgboost [Chen and Guestrin, 2016] with the classifier or regressor and default parameter from the python implementation. We used default parameters and a learning rate of 0.1 and report the accuracies computed with 10-fold validations with 5 randomised repeats.

First, to learn a model of input resistances, we apply the xgboost regressor on the normalized model parameters with Pearson correlation larger than 0.7 (shown in Fig. 10c-h), to prevent any overfitting. If no parameters are correlated enough, we replace the model with the mean value of this parameter. On a new set of models, we then evaluate these models to estimate the input resistance polynomials for both AIS and soma. To prevent our ML model to extrapolate the values, we sub-sample models so that their parameters are between the 10’s and 90’s percentile of the trained set (black lines in Fig. 10c-h).

We train the same regressor model to predict the *ρ* factors with normalized model parameters that have a Pearson correlation larger than 0.4 with the *ρ* factors. The choice for this lower correlation is from the fact that the calibration of the *ρ* factors is based on a coarser scan of the parameter space, hence it is noisier and we do not expect very high correlations. Again, if no parameters are correlated enough, we replace the model with the mean value of this parameter. One the additional models, we again use these ML models to predict their *ρ* factors, but only if their parameters land between the 10’s and 90’s percentile.

Finally, we use the xgboost classifier with the same parameters as previously to estimate electrical model generalisation from their parameter values. Using the Shap feature important analysis [Lundberg and Lee, 2017], we could find the most important parameters to predict the model’s generalisation, shown in Fig. 4c. We then used only models that this classifier predicts as generalisable to test whether our ML calibration produces valid models (see Sec.). We used all normalized model parameters to train this classifier, likely overfitting the results, but without much impact.

### K Increasing morphological variability with neuronal synthesis

In addition to generating many electrical models of a given cell type reproducing experimental data, we can generate morphologies reproducing experimental data for a morphological type. This can be done with neuronal synthesis algorithms trained from the selected population of reconstructions. Several algorithms are available, such as [Luczak, 2006, Koene et al., 2009, Luczak, 2006] but we will use here the more recent, topologically based algorithm of [Kanari et al., 2022]. We generated 100 thick-tufted morphologies for which we adapted the AIS and soma scales for each selected model, and evaluated all pairs of synthesised morphologies and models.

First, in Fig. 9, we confirm from [Kanari et al., 2022] that the experimental morphological variability is well reproduced and in particular the correlations between some main morphometrics. After the evaluation, we find that 94.8% of the pairs of morphologies and models have cost below 5sd and 98.7 have cost below 10sd. In Fig. 11, we show a more detailed analysis of this result, and in particular that only 3 morphologies are responsible for a large part of the costs above 10sd. After further inspections, we found that the surface area of the oblique dendrites is a good predictor of the failure of the morphology on many models. We leave a more detailed analysis of the mechanistic reasons for such a correlation for future works, but it shows that some specific aspects of the branching structure may matter for electrical modelling, even when measured at the soma.

**Figure 11:**
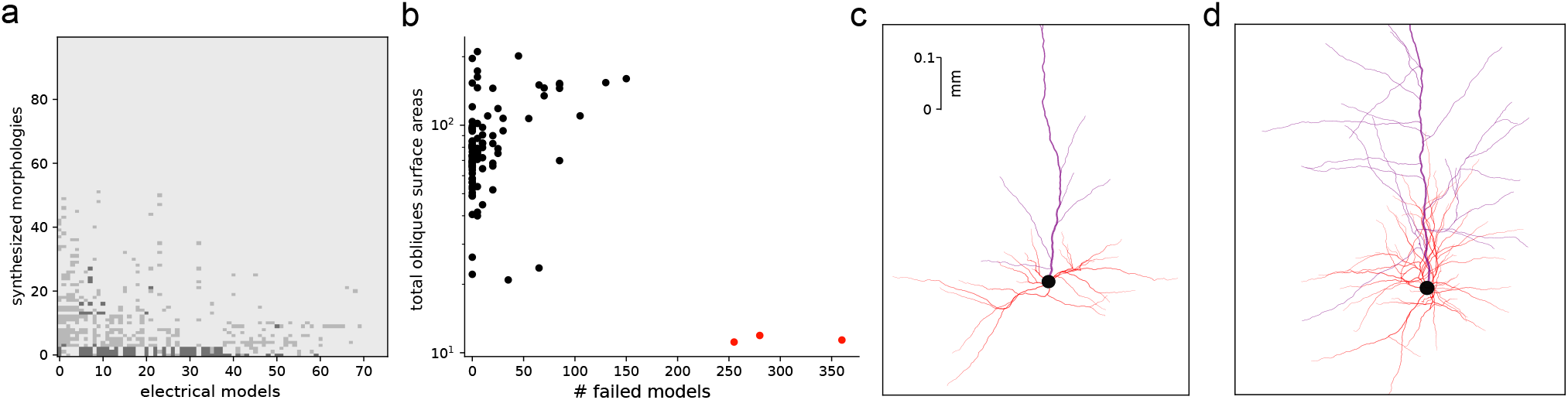
Generalisation with synthesised morphologies. **a** Selection matrix with selected models and synthesise morphologies generated from selected morphologies. Grey pixels correspond to scores above 5 and black for scores above 10. White correspond to scores below 5. **b** For each morphology, we compute the total surface areas of obliques, which is a good predictor of failed morphologies. **c** We show a zoom on the oblique region of a failed morphology. **d** Similar zoom as in **c** but for a working morphology.

### L cNAC electrical model

To show that the MCMC methodology and results also apply to other types of electrical models, we used the continuous non-accommodating (cNAC) electrical model of [Markram et al., 2015, Reva et al., 2022].

In Fig. 12 we show the main results of MCMC and generalisation on Martinotti cells. The exemplar was chosen based on m-types with most reconstructions, with a total of 191 interneurons and 6 mtypes (L23_LBC, L5_MC, L4_LBC, L23_MC, L1_HAC and L4_NBC). The exemplar for MCMC (Fig. 12a) was an L23_MC (Fig. 12b, left) and we illustrate in Fig. 12c,d the generalisation on L5_MC cells.

**Figure 12:**
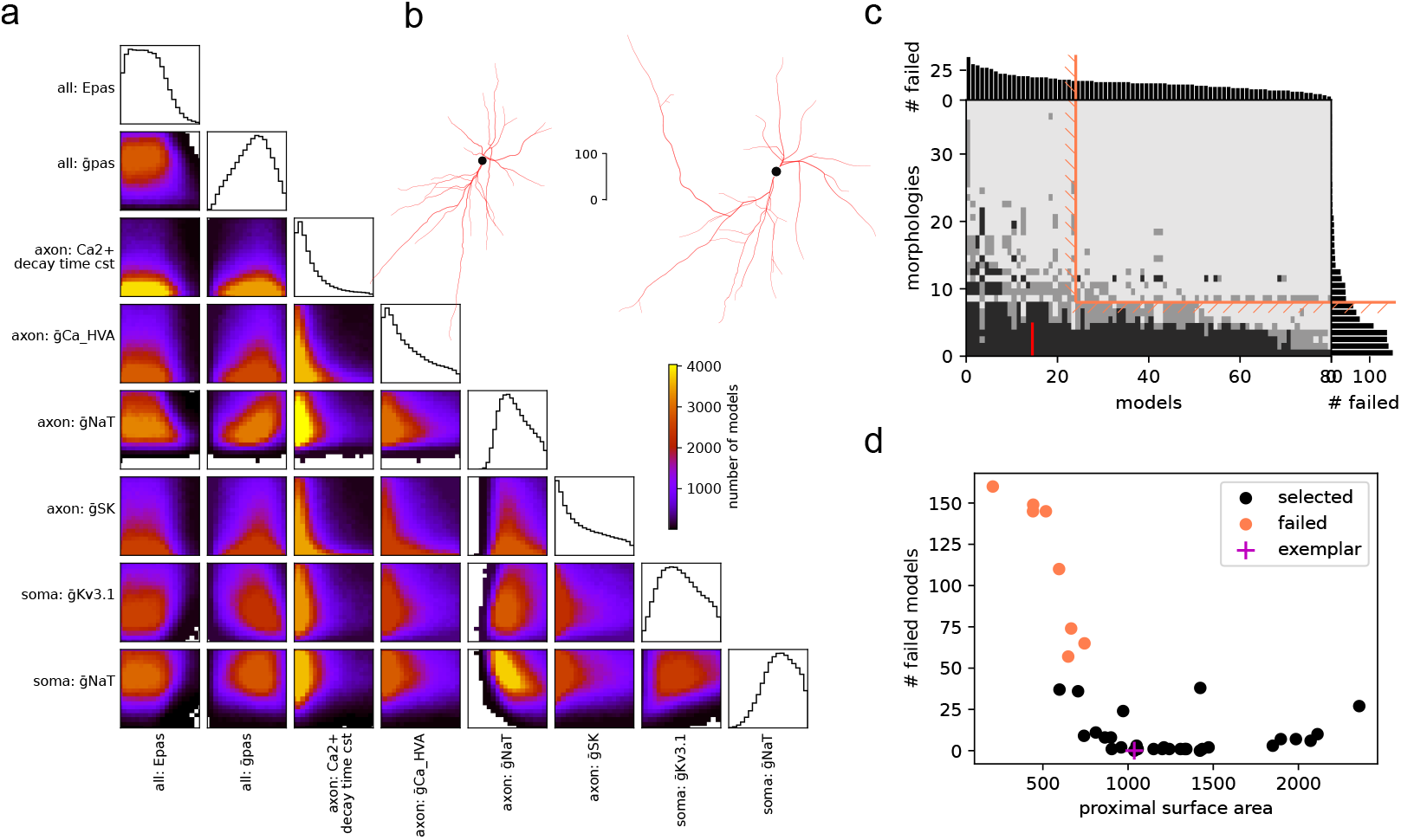
cNAC electrical model. **a** Corner plot of most correlated parameters of an MCMC run on cNAC electrical model based on [Reva et al., 2022]. **b** Exemplar morphologies used for MCMC (left) with a layer 2/3 Martinotti cell and generalisation (right) on layer 5 Martinotti cell. **c** Selection matrix for layer 5 Martinotti cells. **d** Proximal surface areas of cells that are selected and not selected.

**Figure 13:**
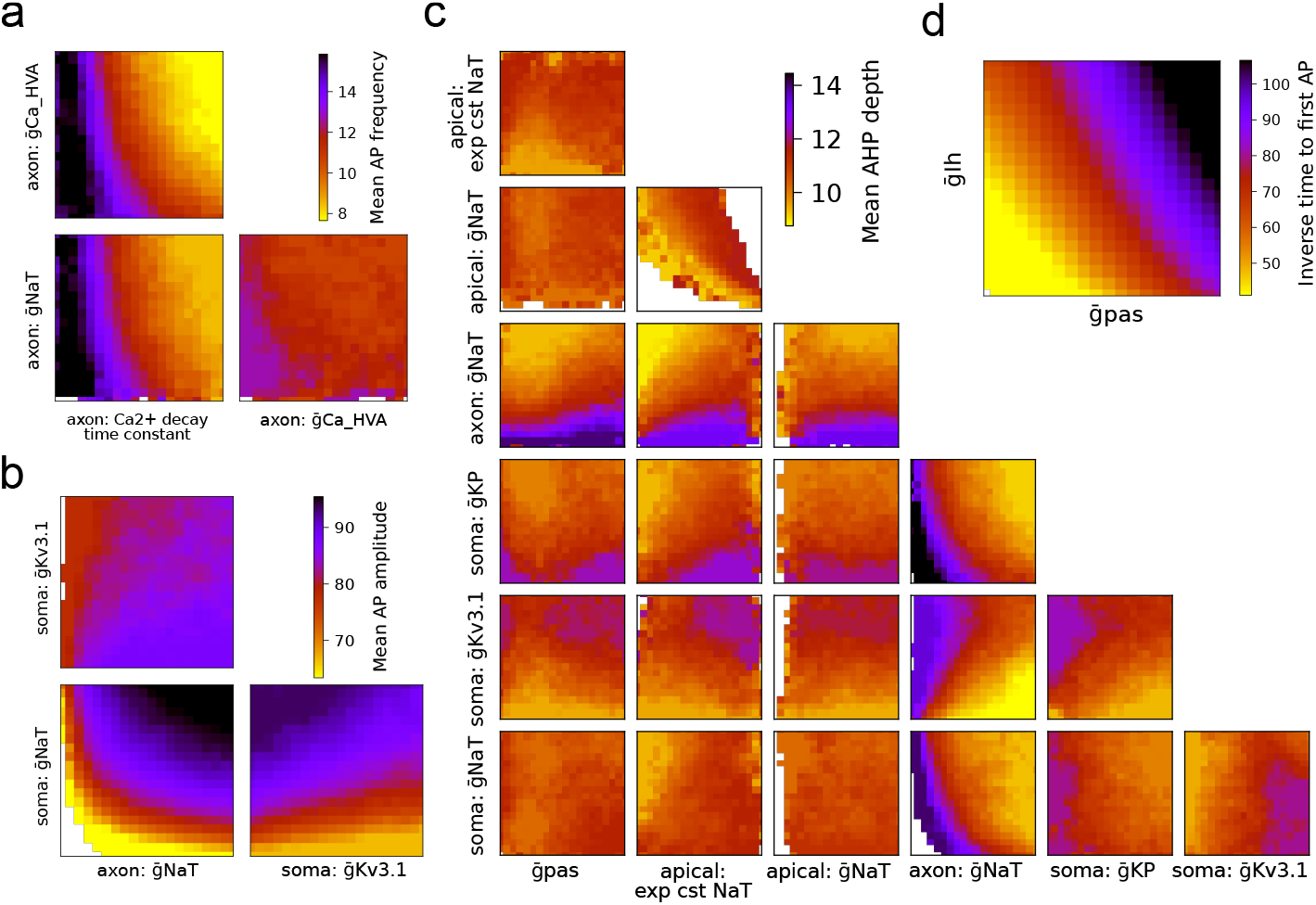
Corner plot with average features values. We plot the average feature values for additional features from Fig. 2f-g. Tee features are: mean AP frequency in **a**, mean AP amplitude in **b**, mean AHP depth in **c** and inverse time to first AP in **d**.

We also applied the generalisation procedure with ML models of *ρ* factors and AIS/soma input resistance to obtain 87.1% of the 7620 evaluated pairs with a score below 5sd, and 98.7% then were able to spike.

As seen from the corner plot, we did not attempt to adjust the bounds of the parameter space precisely in this example, but more work could be done to refine this interneuron model to perform comparisons with other electrical types.

### M Additional figures

**Figure 14:**
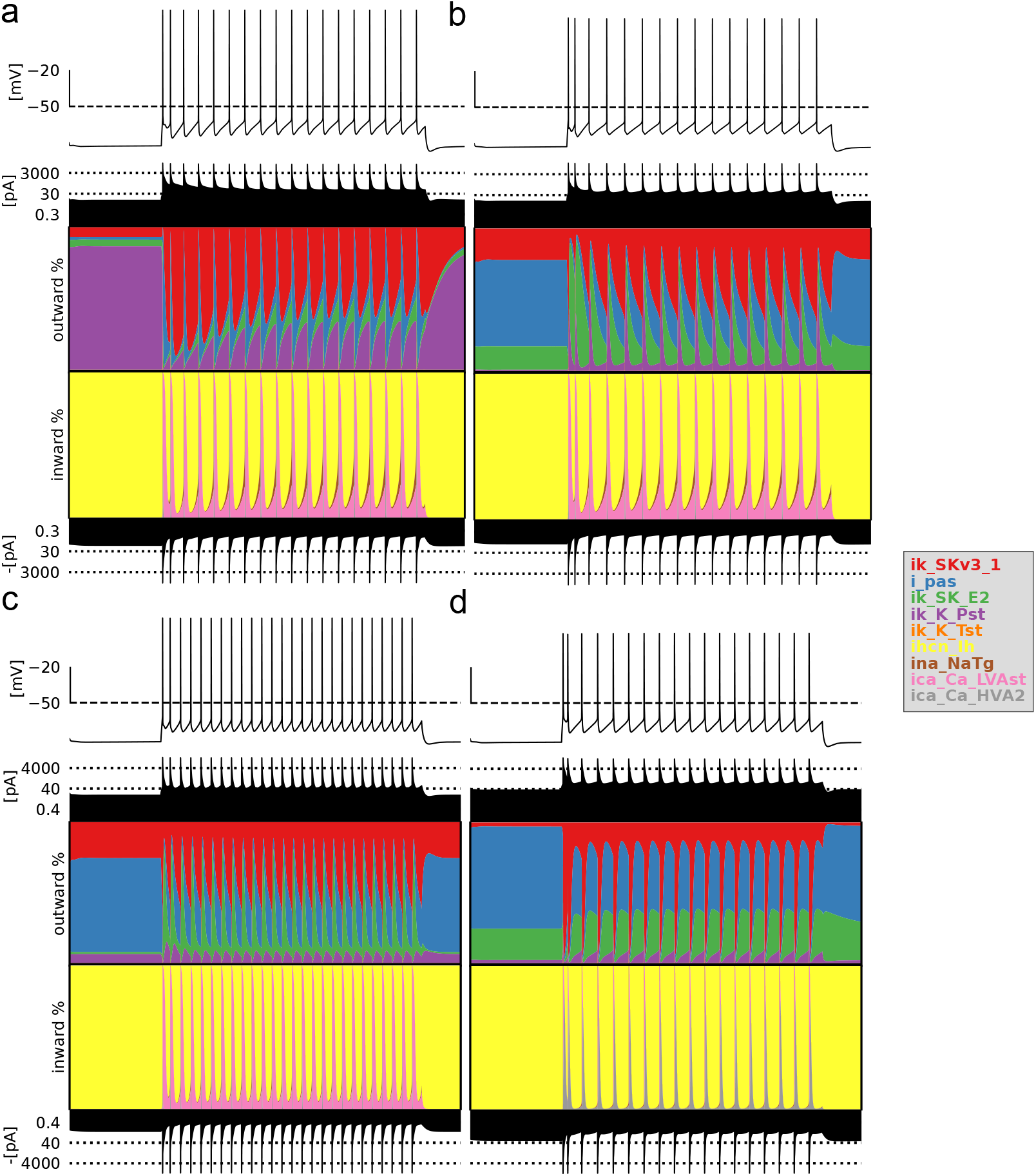
Currentscape of four MCMC models. Currentscape are made following [Alonso and Marder,2019] and represent the various inwards and outwards currents present in the cell during activity with the following models from Fig. 2: blue in **a**, red in **b**, magenta in **c** and green in **d**.

**Figure 15:**
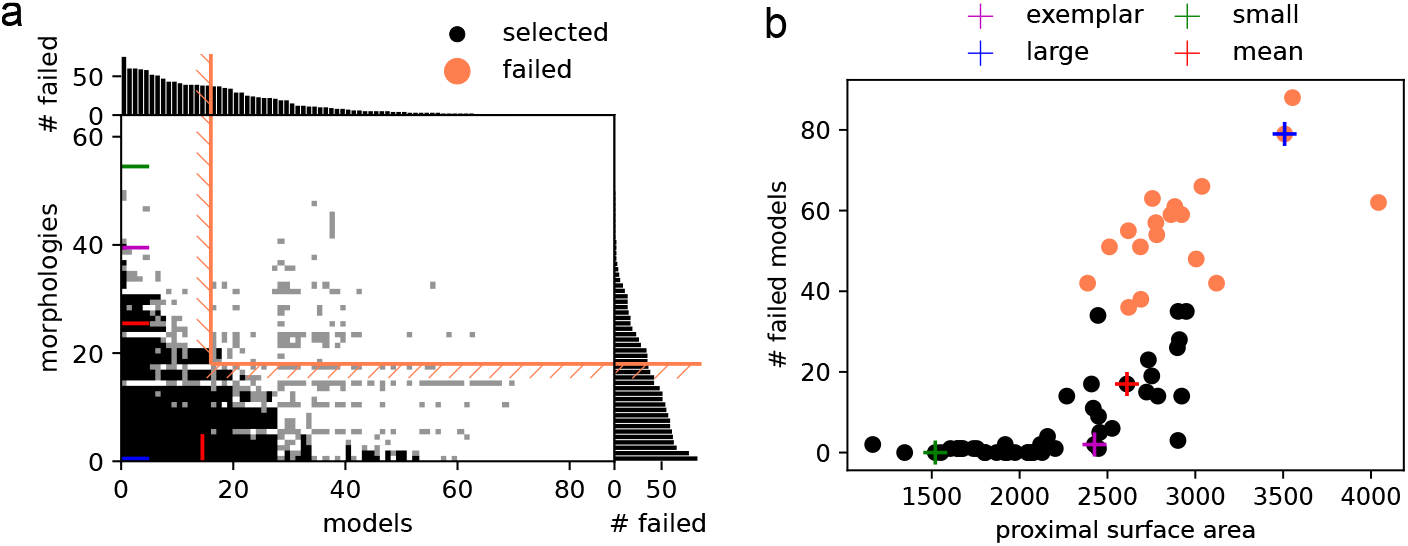
Selection of model and morphologies without AIS/soma adaptation. Same panels as Fig. 4a-b, but without adaptation of AIS/soma.

